# QuantaMind MD enables protein modeling with ab initio accuracy

**DOI:** 10.1101/2025.09.01.673405

**Authors:** Song Xia, Deqiang Zhang, Xu Shang, Jinbo Xu

**Affiliations:** MoleculeMind

## Abstract

Accurate biomolecular simulations are essential for understanding chemical processes and advancing applications such as protein engineering and drug design. While quantum mechanical (QM) methods can provide chemical accuracy, their computational cost limits their scalability. Machine learning force fields (MLFFs) offer a promising alternative by achieving similar accuracy with vastly improved efficiency, but their effectiveness is often constrained by limited conformational and chemical diversity in training. We introduce QuantaMind, a robust MLFF workflow designed to extend chemical and conformational coverage, particularly in transition-state regions. QuantaMind enables quantum-accurate molecular dynamics simulations, successfully capturing complex phenomena such as proton diffusion, water dissociation, and acid-base neutralization. We show that QuantaMind can be applied to large biomolecular systems, enabling accurate protein structure optimization and improving the prediction of residue contacts and hydrogen bonds. A pocket-centric simulation strategy further allows QuantaMind to efficiently model protein-ligand interactions with high structural accuracy. These results establish QuantaMind as a versatile and scalable tool for atomistic simulations at ab initio accuracy.

## Introduction

Molecular dynamics (MD) simulations are crucial computational biomolecular modeling, offering insights into the structure, dynamics, and function of complex molecular systems.^1–3^ In a typical MD simulation, atomic trajectories are generated by computing interatomic forces at each time step and integrating Newton’s equations of motion.^4^ In principle, the most accurate way of calculating the atomic force involves solving the Schrodinger equation (SE).^5^ Unfortunately, analytical solutions are only available for systems with few degrees of freedom. Numerical solutions such as high-accurate quantum mechanics (QM) calculations impose prohibitive computational cost.^6,7^

Classical force fields (FFs) use empirical functions to approximate the energies and atomic forces of the system.^8,9^ While classical FFs enable simulations of systems with thousands of atoms, their accuracy is ultimately limited by simplifications and fixed parameters that fail to capture complex electronic effects, especially in reactive environments.

Machine learning force fields (MLFFs) circumvent the task of solving the SE by directly learning the mapping from the geometry to the energy, forces, and other properties of the system through supervised learning,^10–12^ offering orders of magnitudes of speed up. Recently, a growing number of MLFFs have demonstrated promise in chemical and biomolecular domains.^13–17^ However, most of the MLFFs for biomolecular simulations are trained on equilibrium or near-equilibrium conformations sampled from classical FF simulations. As a result, their accuracy declines when applied to systems outside the distribution of the training set, such as those involving chemical reactions or diverse elemental compositions.^18^

In this study, we introduce QuantaMind, a deep learning-based MLFF workflow designed to support a diverse range of chemical environments and biomolecular systems. To address the limitations of prior models, we include transition state data to capture chemically relevant configurations at non-standard bond lengths and angles. In addition, we construct a dedicated phosphorus-containing dataset to expand element diversity in the training regime. We demonstrate that QuantaMind can accurately simulate bond formation and breaking events and achieves good agreement with experimental data. On a larger scale, QuantaMind supports protein structure refinement and protein-ligand interaction prediction via efficient, scalable simulations. This work establishes QuantaMind as a general-purpose AI platform for atomistic modeling with ab initio-level accuracy.

## Results and discussion

### Accurate energy and force prediction with QuantaMind

Figure 1A presents an overview of the QuantaMind training protocol. The cycle begins with a biomolecular simulation of the system of interest. Classical molecular dynamics (MD) simulations are used to model large biomolecular systems. Transition state optimization using high-level quantum mechanics (QM) methods is frequently employed to study enzymatic reactions, which involve the breaking and formation of chemical bonds. The QuantaMind framework developed in this work can also be applied to simulate large biomolecular systems with ab initio accuracy, potentially contributing to this cycle.

**Figure 1:**
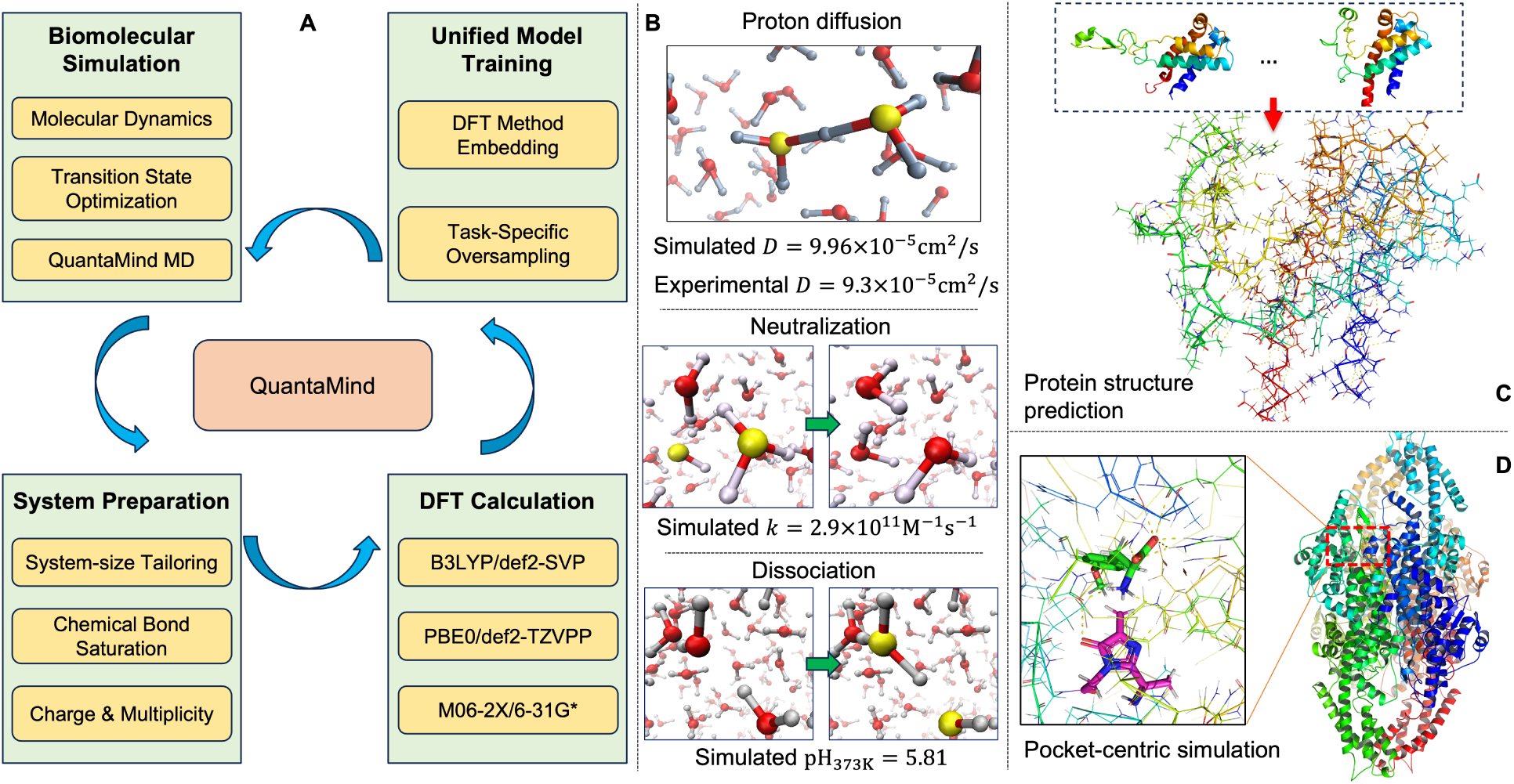
Development and application of QuantaMind. (**A**) The overview of QuantaMind training protocol. (**B**) Simulation of proton diffusion, acid-base neutralization and spontaneous water dissociation. (**C**) Protein dynamic structure prediction at atomic level with ab initio accuracy. (**D**) Pocket-centric simulation for protein-ligand interaction prediction.

After the biomolecular simulation, the next step involves preparing the system for QM calculations. Due to the high computational cost of high-level QM methods, the system is divided into fragments, each containing fewer than 300 atoms. After fragmenting the system, bond saturation is performed to maintain proper chemical validity where bonds have been severed. The total charge and multiplicity of each fragment are determined based on chemical knowledge in preparation for QM calculations.

QM properties-including energy, forces, and dipole moments-are calculated using Density Functional Theory (DFT) methods. ^19^ To balance accuracy with computational efficiency, the choice of DFT functionals and basis sets depends on the specific system and type of calculation.^20^ As a result, different datasets may have DFT properties computed using different combinations of functionals and basis sets. To address this variability, we introduce a DFT method embedding strategy that encodes the functional and basis set information, enabling the model to be aware of the source of each label. Additionally, an oversampling strategy is employed during training to ensure the model has an equal opportunity to learn from examples across datasets.^21^ This is particularly important when training on datasets of varying sizes.

The trained QuantaMind model can be used for a broad range of protein design applications, including accurate simulation of protons in aqueous solution (Figure 1B and Section Ab initio simulation of proton diffusion, acid-base neutralization and water dissociation), protein dynamic structure prediction at atomic level with ab initio accuracy (Figure 1C and Section Protein dynamic structure prediction with ab initio accuracy) and pocket-centric simulation for protein-ligand interaction prediction (Figure 1D and Section Pocket-centric simulation for protein-ligand interaction prediction).

Following the training protocol illustrated in Figure 1A, we developed four datasets as part of this work. As shown in Figure 2A, dataset P1 highlights phosphorus atoms, which are commonly found in biomolecular systems but are not covered by previously developed QM datasets such as GEMS^16^ and AI2BMD.^17^ Dataset TS1 focuses on the non-canonical amino acid 3,5-dihydro-5-methylidene-4H-imidazol-4-one and its associated enzymatic reaction. Dataset TS2 emphasizes a metal ion and its role in a catalytic reaction. Dataset TS3 features an epoxy group and the hydrolysis reaction catalyzed by a specific enzyme. Details about these two datasets are covered in Section Phosphorus data set construction and Section Transition state dataset construction.

**Figure 2:**
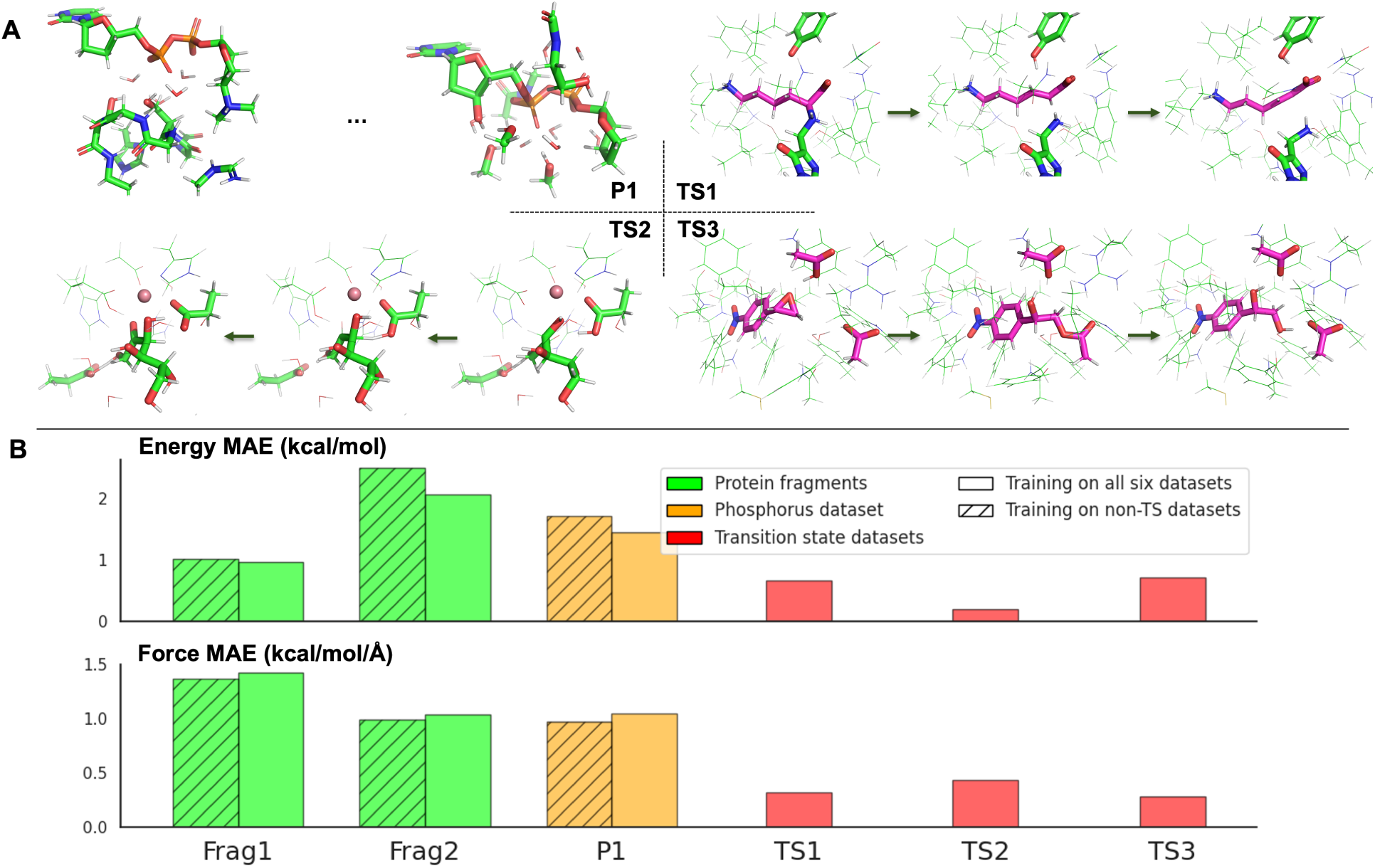
Model achieves state-of-the-art performance on protein fragments, our created phosphorus dataset and our transition state dataset. (**A**) Data sets created in this work. P1 is the phosphorus data set (see Section Phosphorus data set construction). TS1, TS2 and TS3 are the transition state data sets.(see Section Transition state dataset construction) (**B**) Model performance on two public data sets and four data sets created in this work.

In addition to the four datasets developed in this work, we also incorporate the General Protein Fragments and Crambin datasets-referred to as Frag1 and Frag2, respectively-from the GEMS^16^ study to support model training. Figure 2B presents the model performance in terms of mean absolute error (MAE) for energy and force predictions across the test sets of all six datasets. Although the transition state datasets were generated using different combinations of functionals and basis sets, incorporating them into the training set does not significantly degrade model performance. This demonstrates the capability of QuantaMind to learn a unified molecular representation for predicting DFT-level properties across diverse data sources.

### Ab initio simulation of proton diffusion, acid-base neutralization and water dissociation

Conventional molecular mechanics simulations are limited in their ability to model chemical bond breaking and formation. In contrast, QuantaMind, a deep learning model trained from QM data, does not suffer from this constraint. To assess its capabilities, we employed QuantaMind to simulate a single excess proton in a water box (Figure 3A). Over a 1 nanosecond (ns) trajectory, we observed multiple proton transfer events between water molecules (Figure 3B).

**Figure 3:**
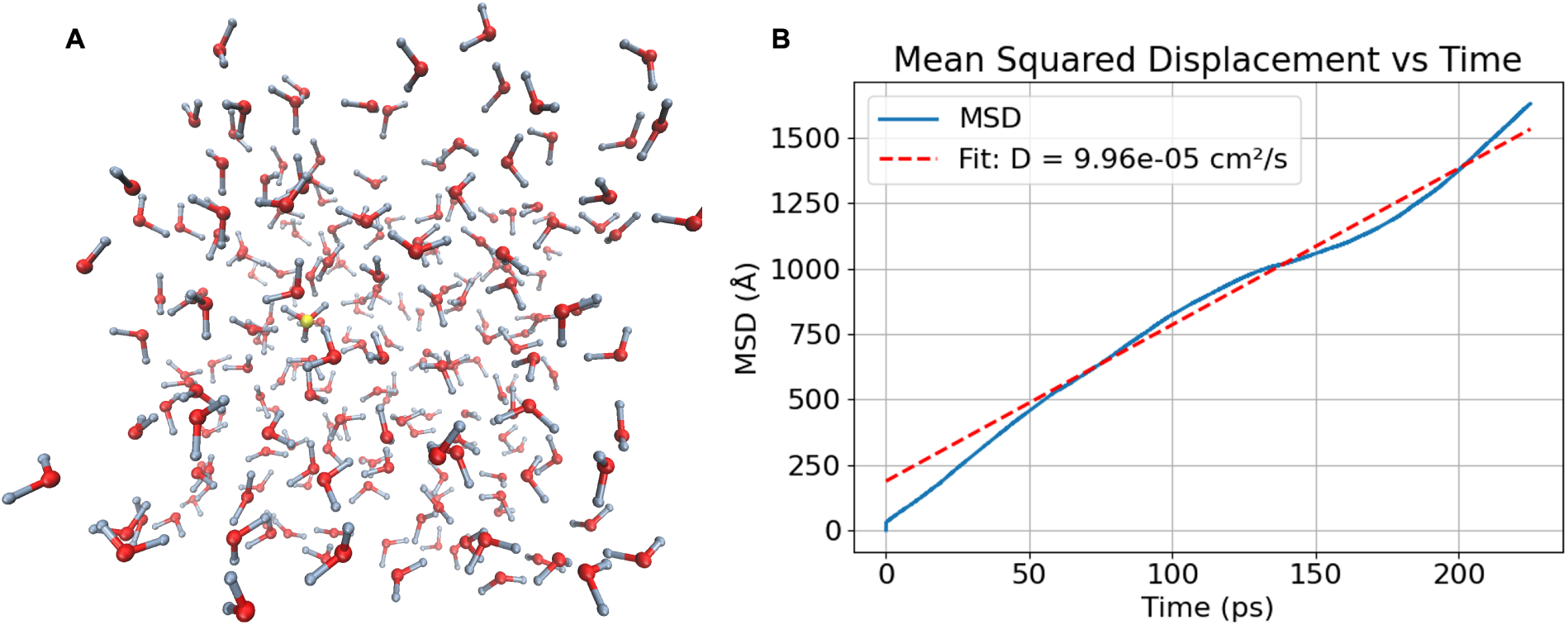
Simulation of a proton in a water box with QuantaMind. (**A**) Visualization of the simulation system, with the hydronium ion highlighted in yellow. (**B**) Mean squared displacement (MSD) versus time and fitted diffusion constant.

To quantify the proton’s mobility, we estimated the diffusion coefficient using the Einstein relation:^22,23^

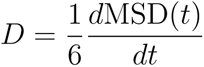

where MSD(*t*) denotes the mean squared displacement as a function of time. Figure 3B shows the linear dependence of MSD on time, from which we derive a diffusion coefficient of 9.96 × 10*^−^*^5^cm^2^*/*s. For comparison, the experimentally measured diffusion coefficient of a proton in water is approximately 9.3 × 10*^−^*^5^cm^2^*/*s. Notably, QuantaMind produces this estimate without specific fine-tuning on proton transfer data, demonstrating its robustness and potential to capture key dynamical behavior consistent with experimental observations. We further evaluated QuantaMind’s ability by modeling the acid-base neutralization reaction in water. Using the same simulation box as in the previous proton diffusion study, we replaced one water molecule with a hydroxide ion, positioned approximately 10 Å away from an existing hydronium ion. Five independent simulations were conducted with randomized initial velocities to capture the stochastic nature of molecular dynamics.

Figure 4A shows the distance over time between the hydronium and hydroxide ions in all five simulation runs. In several cases (e.g., runs 1, 2, 3, and 5), we observe a sharp drop in distance to near zero around 12 ps, corresponding to a neutralization event. Following this event, QuantaMind no longer detects ionic species, indicating the formation of two neutral water molecules. In run 4, however, the ions do not encounter each other during the 50 ps simulation window.

**Figure 4:**
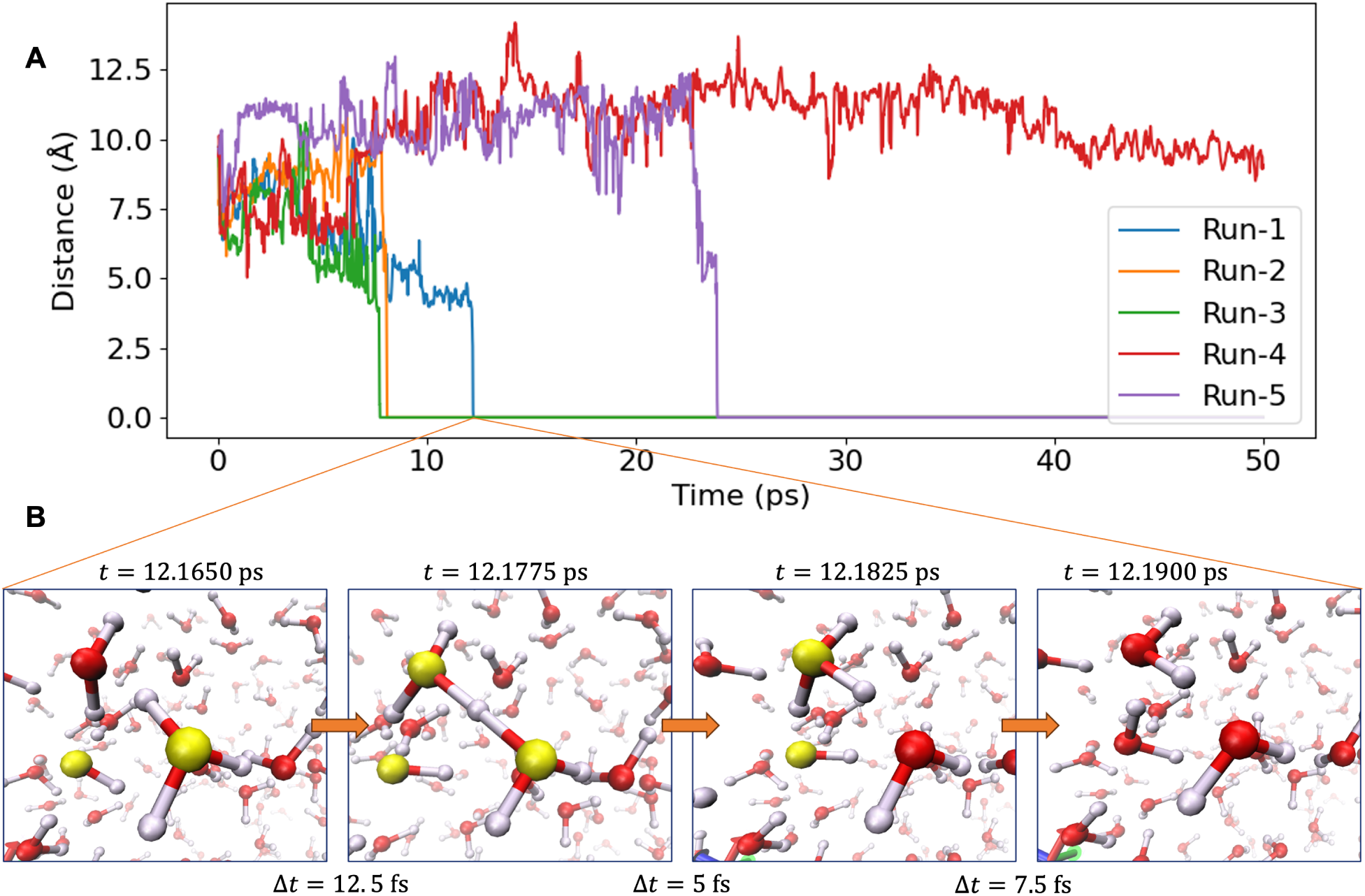
Simulation of an acid-base neutralization reaction in aqueous solution. (**A**) Distance between the hydronium ion and hydroxide ion over time from five independent simulation runs. (**B**) Visualization of the neutralization event at 12.1650 ps in run 1. The oxygen atoms in the hydronium and hydroxide ions are highlighted in yellow.

Figure 4B depicts the neutralization event for run 1 at 12.1650 ps. The visualized trajectory reveals a rapid proton transfer from the hydronium ion to a neighboring water molecule (first three frames), immediately followed by the neutralization of the hydroxide ion, forming two water molecules. The whole process take places within 25 fs.

To estimate the rate constant of the neutralization event, we use the following formula:

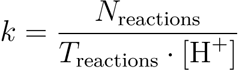

where *N*_reactions_ = 1 is the number of reactions observed per simulation, *T*_reactions_ is the average time required for a neutralization event to occur, and [H^+^] is the proton concentration. The reaction times for the five independent trajectories are:

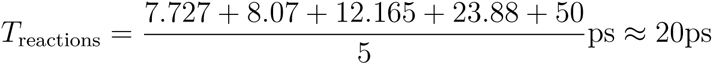

Here, 50 ps is used for run 4 as a conservative estimator since the reaction did not occur in that window.

The proton concentration is calculated as:

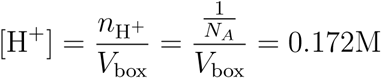

where *N_A_* = 6.022 × 10^23^*mol^−^*^1^ is the Avogadro constant. *V*_box_ = 21.18 · 21.47 · 21.18 Å^3^ is the volumne of our simulation box. Combining these values, we estimate the reaction rate constant:

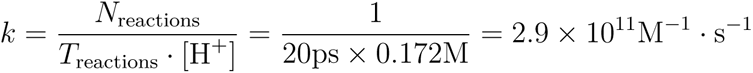

It is worth noting that acid-base neutralization reactions are expected to proceed significantly faster than typical diffusion-controlled processes. This is largely due to the fact that both protons and hydroxide ions can propagate rapidly through the hydrogen-bond network of water. In addition, electrostatic steering between the oppositely charged ions enhances the likelihood of their recombination. While conventional diffusion-controlled reactions generally occur at rates between 10^9^ and 10^10^ M*^−^*^1^ · s*^−^*^1^,^24^ our calculations indicate that acid-base recombination in water reaches rates on the order of 10^11^ M*^−^*^1^ · s*^−^*^1^, confirming that this process is noticeably faster than typical molecular diffusion limits.

Since acid-base neutralization occurs instantaneous for most experimental techniques, our simulations using QuantaMind provide valuable insights by capturing the explicit steps of the neutralization process in atomistic detail and enabling an estimation of the reaction rate constant.

In principle, QuantaMind is capable of simulating the reverse reaction of acid-base neutralization–specifically, the autoionization of water into hydronium (H3O+) and hydroxide (OH-) ions. However, due to the extremely low probability of such events under standard conditions, capturing a spontaneous water dissociation event via molecular simulation is computationally demanding. To increase the likelihood of observing such an event, we conducted two independent 0.5 ns simulations of a neutral water box at an elevated temperature of 373 K.

Figure 5 summarizes the results of the simulations. At each frame, we quantify the number of hydronium and hydroxide ions by evaluating the distances between oxygen and hydrogen atoms, based on geometric criteria consistent with proton transfer. As expected, for the majority of the simulation time, no ions are detected, indicating that all water molecules remain in their neutral form. However, we observe a transient increase in the number of detected ion pairs at 462.9 ps in run 1 and 347.6 ps in run 2, suggesting the occurrence of a water dissociation event.

**Figure 5:**
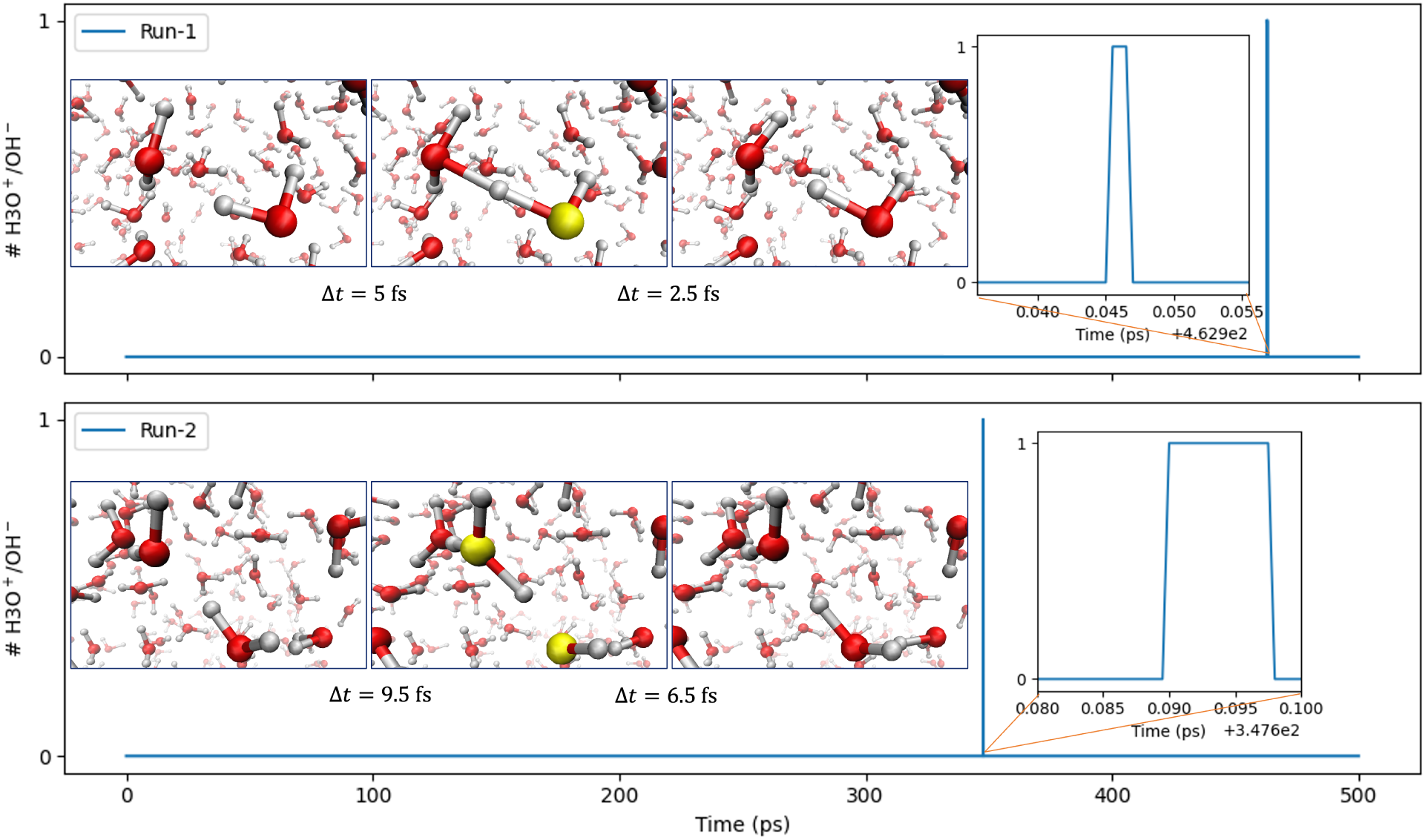
Sampling of water dissociation events at 373 K. The number of hydronium and hydroxide ions detected over time is shown for two independent simulation runs.

Closer inspection of the molecular trajectories confirms these events. In run 1, we observe a proton from one water molecule being transiently shared with a neighboring water, resulting in a short-lived hydronium-like species characterized by an oxygen atom bonded to three protons. The proton rapidly returns to its original donor, re-forming two neutral water molecules. In run 2, a proton transfer occurs in which a hydrogen atom jumps from one water molecule to another, forming a hydronium and a corresponding hydroxide ion. This ion pair persists for approximately 6.5 fs before the proton transfers back to the original molecule, thereby reestablishing a neutral water configuration.

Across our two independent 500 ps simulations, water dissociation was observed for a total of 9.0 fs (2.5 fs in run 1 and 6.5 fs in run 2). The probability of capturing such a dissociation event in our system is estimated as:

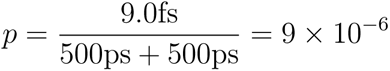

Given that each observed dissociation event corresponds to a proton (or hydroxide ion) concentration of 0.172 M, as discussed above, the estimated equilibrium concentration of

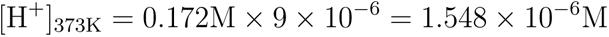

Accordingly, the estimated pH at 373 K is:

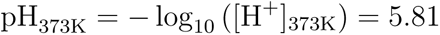

This value is in reasonable agreement with the experimentally reported pH of 6.14 for pure water at 373 K, supporting the accuracy of QuantaMind in modeling rare but fundamental chemical fluctuations.

We acknowledge that pH is an equilibrium property, and a more rigorous calculation would require significantly larger simulation volumes and longer time scales. However, capturing such ultrafast events requires high temporal resolution-our trajectories were recorded every 0.5 fs, resulting in approximately 25 GB of data per 500 ps simulation of 216 water molecules. Extending the simulation length would substantially increase both computational costs and storage demands.

### Protein dynamic structure prediction with ab initio accuracy

We apply QuantaMind to protein structure prediction, with a particular focus on side-chain contact and hydrogen bond prediction. The overall workflow is illustrated in Figure 6A. Starting from a protein sequence, backbone sampling tools are used to generate a variety of possible backbone conformations (Route 1). In this work, we use BioEmu^25^ as the backbone sampling tool; however, alternative tools such as Str2Str,^26^ ESMDiff^27^ and AlphaFlow^28^ are also applicable. Side-chain packing is then performed to build side-chain structures based on the generated backbones. We employ HPacker^29^ in this study, although other tools such as AttnPacker^30^ and RosettaPacker^31^ can also be integrated into the workflow. Following side-chain packing, a quick structure relaxation is carried out using Rosetta FastRelax.^31^ Finally, QuantaMind is used to optimize the resulting protein structure.

**Figure 6:**
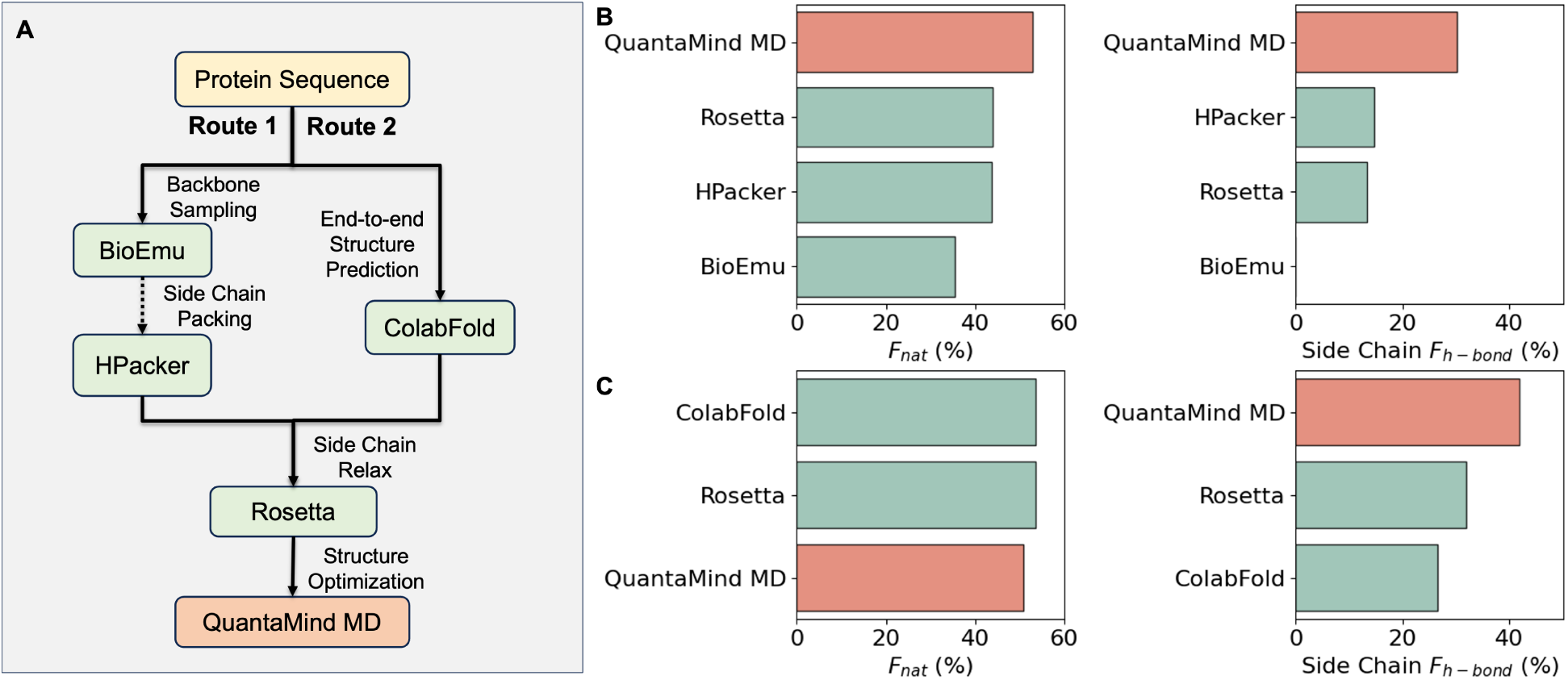
Evaluation of protein structure prediction tasks on a benchmark of 17 protein examples. (**A**) Two protein structure prediction workflows. In route 1, first the backbone structures are sampled and then the side chain structures are predicted. In route 2, the protein structure is predicted directly. A quick side chain relaxation is performed to solve side chain clashes. QuantaMind MD is then employed to optimize the protein structure. (**B**) Model performance in terms of *F*_nat_ and side chain *F*_h-bond_ following route 1. (**C**) Model performance in terms of *F*_nat_ and side chain *F*_h-bond_ following route 2.

We evaluate model performance using two metrics. The fraction of native contacts (*F*_nat_) measures the proportion of native residue–residue contacts preserved in the predicted structure. Side-chain hydrogen bond recovery (Side Chain *F*_h-bond_) quantifies the proportion of native hydrogen bonds retained in the predicted structure. Figure 6B summarizes the average model performance on our benchmark set of 17 protein examples (see Section Protein evaluation benchmark and metrics). We observe improvements in both *F*_nat_ and *F*_h-bond_ following QuantaMind optimization.

Alternatively, end-to-end structure prediction models, such as those predicting both backbone and sidechain structures, can be seamlessly integrated into our workflow (Figure 6A, Route 2). We use ColabFold^32–34^ for the structure prediction, followed by structure relaxation with Rosetta FastRelax and QuantaMind. Figure 6C summarizes the average model performance on our benchmark set. We observe comparable performance in *F*_nat_ prediction and an improvement in *F*_h-bond_ following QuantaMind optimization.

### Pocket-centric simulation for protein-ligand interaction prediction

Predicting protein-ligand interactions is critically important in both drug discovery and enzyme design. One of the main challenges in using QuantaMind to simulate protein-ligand systems lies in the chemical diversity of small molecules, which far exceeds that of amino acids. To evaluate QuantaMind’s performance on small molecules, we conducted an energy optimization task on a benchmark set of 1,500 small molecules. We then computed the DFT energies of each molecule before and after optimization using accurate QM methods. Figure S1A presents the distribution of energy differences across the dataset. Notably, most optimizations resulted in lower-energy conformations, indicating the effectiveness of QuantaMind in refining small molecule geometries.

Given that the model is primarily trained on biomolecular systems composed of amino acids, we assess how well a small molecule fits within the model’s learned chemical space by computing its maximum structural similarity to the 20 standard amino acids. We use this similarity score as a proxy for model uncertainty. Figure S1B illustrates a scatter plot of DFT energy differences as a function of molecular similarity. Figures S1C and S1D show the distribution of energy differences and the success rate of optimizations across different confidence intervals, respectively. An optimization is considered successful if the post-optimization DFT energy is lower than the pre-optimization value (i.e., energy difference < 0).

For protein-ligand interactions, inspired by QM/MM approaches that focus computational efforts on the local binding environment, we implemented a pocket-centric simulation algorithm. This approach restricts simulations to residues within a specified distance from ligand atoms, significantly reducing computation for large proteins by avoiding unnecessary calculations far from the binding site.

Figure 7A illustrates a typical protein-ligand structure optimization workflow. Starting from AlphaFold3^36^ predicted structures on the PoseBuster dataset,^35^ we apply Rosetta’s FastRelax^31^ protocol for sidechain relaxation. Subsequently, the structure undergoes pocket-centric optimization with QuantaMind. Figure 7B summarizes the model performance on the PoseBuster dataset. Based on the analysis in Figure S1D, we exclude entries with ligand similarity scores below 0.44. Across the remaining dataset, QuantaMind achieves improvement across all intra- and intermolecular validation metrics. However, no improvement is observed for ligand RMSD. To further evaluate model performance, we measure hydrogen bond recovery, denoted as F_h-bond_, which quantifies the fraction of protein-ligand hydrogen bonds present in the optimized structure relative to the crystal structure. We observe a marked increase in F_h-bond_ for structures optimized by QuantaMind compared to those optimized by Rosetta or derived from AlphaFold3 predictions.

**Figure 7:**
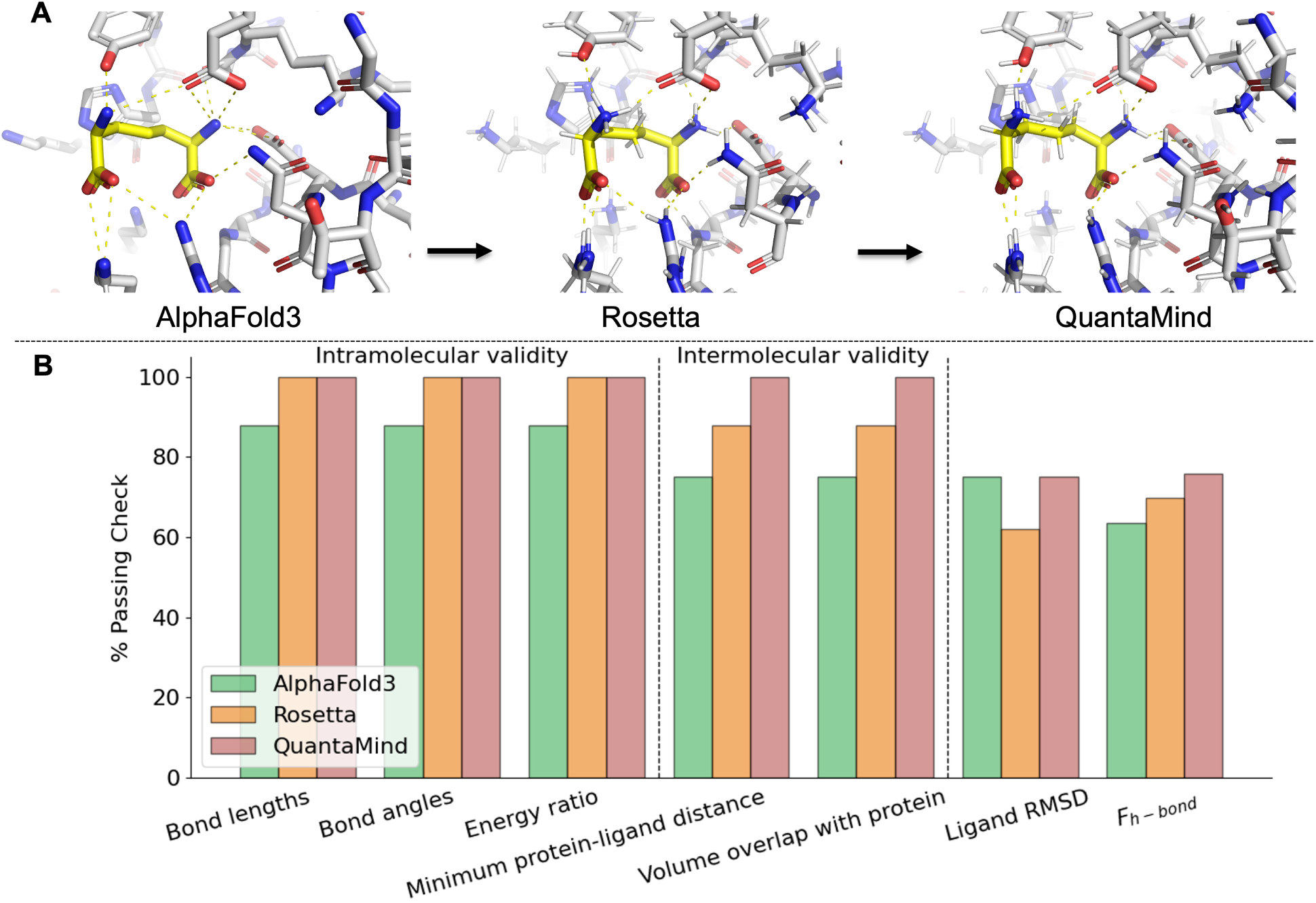
QuantaMind delivers accurate protein-ligand interaction predictions through pocket-centric simulation. **A.** Example protein-ligand optimization using the QuantaMind workflow. **B.** Model performance on the PoseBuster^35^ dataset after filtering for ligands with similarity scores greater than 0.44.

## Discussion

In this work, we have developed the QuantaMind workflow–a neural network-based platform that achieves state-of-the-art performance on molecular property prediction tasks, including energy and force estimation. Notably, molecular dynamics simulations driven by QuantaMind accurately reproduce the experimental diffusion coefficient for protons. Beyond proton diffusion, the framework also enables simulations of acid-base neutralization and water dissociation events.

At larger scales, we demonstrate that QuantaMind can be effectively applied to protein structure optimization, yielding improvements in both residue-residue contacts and hydrogen bond recovery. Furthermore, by implementing a pocket-centric simulation paradigm, we significantly reduce computational overhead while enhancing the precision of structural predictions in ligand-binding sites.

Despite its strengths, a key current limitation of the QuantaMind model is its relatively narrow training regime on small molecules. Exploring the vast chemical space has been an active area of research,^37–40^ and many comprehensive QM datasets have been published in recent years.^16,17,41–44^ Incorporating such datasets into the training pipeline could improve the model’s generalizability across diverse chemistries, particularly benefiting protein-ligand simulations.

For transition state optimization, we have evaluated the model on three hand-picked systems as a proof-of-concept. While these systems provide illustrative use cases, we recognize the availability of benchmark datasets specifically curated for TS tasks, which span broader chemical and conformational spaces.^45,46^ Integrating these datasets in future work can enhance QuantaMind’s robustness and accuracy in modeling reaction pathways. Machine-learned force fields for TS optimization remain an active research frontier,^46–50^ predominantly focused on small molecules. We believe QuantaMind offers a promising avenue to extend these techniques into biomolecular regimes, paving the way for AI-accelerated enzyme and catalyst designs with QM accuracy.

Finally, the pocket-centric simulation strategy we introduced can be further generalized into an interface-centric paradigm. This extension would be particularly valuable for modeling protein-protein interactions, such as those involved in antibody-antigen binding or complex assembly. By enabling focused, high-accuracy simulations at molecular interfaces, QuantaMind could empower next-generation predictive tools for biotherapeutics and structural biology.

## Materials and Methods

### Phosphorus data set construction

The phosphorus dataset is derived from the spinosyn forosaminyltransferase enzyme, which catalyzes the glycosylation reaction between 17-pseudoaglycone and TDP-D-forosamine to form spinosyn and TDP.^51^ The initial protein structure was obtained from the crystal structure of the enzyme available in the RCSB PDB under the accession code 4LEI. To resolve missing regions in the crystal structure, we incorporated loop predictions from the AlphaFold2 model. Subsequently, we employed AutoDock Vina^52^ to dock both reactants– 17-pseudoaglycone and TDP-D-forosamine–into the enzyme’s putative binding pocket. The resulting protein-ligand complex was then subjected to classical molecular dynamics (MD) simulations in explicit solvent to sample conformational space.

After simulation, we extracted 1,000 frames of the protein-ligand complex. Since each full system contains over 50,000 atoms–far beyond the feasibility of DFT calculations–we developed a system-tailoring algorithm to reduce the model size to fewer than 300 atoms, suitable for QM calculations. The algorithm includes the following steps:

1. Identify a point-of-interest (POI) atom. In this context, the POI is either a phosphorus atom in TDP-D-forosamine or a randomly selected atom within 5 Å of the phosphorus atom.
2. Select all atoms within a 5 Å radius of the POI. Chemical bonds are truncated at the boundary of this sphere.
3. Saturate truncated bonds: for each single bond cut at the boundary, if the retained fragment includes a heavy atom (non-hydrogen), replace the missing bonded atom with a hydrogen. The hydrogen position is determined based on the original bond orientation and standard single-bond lengths.
4. If a cut bond is not a single bond (e.g., a double or aromatic bond) or the retained fragment is a single hydrogen, retain the bonded atom and expand the boundary. The algorithm recursively evaluates new terminal bonds until all cut bonds meet the single bond saturation criterion.

Using this approach, we extracted three chemically valid substructures from each simulation frame, resulting in a total of 3,000 structures. Each structure was subjected to DFT single point energy and force evaluation using the Psi4 software package^53^ at the PBE0/def2-TZVPP+MBD level of theory.^54,55^ After filtering for successful calculations, a total of 2,476 optimized molecules were retained, comprising the final phosphorus dataset.

### Transition state dataset construction

The transition state (TS) datasets used in this work were derived from quantum mechanical (QM) cluster simulations.^56^ Three datasets-TS1, TS2, and TS3-were created to represent distinct enzyme-catalyzed reaction pathways and broaden the coverage of transition-state configurations. Importantly, we note that these datasets were originally generated as part of independent research projects and not specifically designed for the purpose of training QuantaMind.

TS1 focuses on the enzymatic activity of phenylalanine ammonia-lyase (PAL),^57^ which catalyzes the conversion of L-phenylalanine into trans-cinnamic acid and ammonia. In addition to L-phenylalanine, we explored alternate substrates, including L-tyrosine and L-lysine, to expand the chemical diversity. Crystal structures with PDB IDs 1W27, 1Y2M, 6RGS, 6HQF, 6F6T and 6H2O were used as templates, and ligand conformations were predicted using AutoDock Vina. ^52^

TS2 targets D-psicose 3-epimerase,^58^ initiating from crystal structures with PDB IDs 3VNK, 3VNJ, and 8XIV. Similarly, TS3 explores epoxide hydrolases and their associated enzymatic reactions,^59^ using structures with PDB IDs 5AIF, 5AIG, 5AIH, and 5AII. Native reactant conformations that are not present in the crystal structures, such as (p-Nitrophenyl)oxirane, were predicted using AutoDock Vina.

For all three systems, transition state searching and structure optimization were conducted using the Gaussian 16 package^60^ at the B3LYP/def2-SVP level of theory. Each frame obtained during the transition state optimization, along with the corresponding single-point energy and atomic forces, was included in the dataset. TS1 comprises 728 entries, TS2 contains 2,045 entries, and TS3 comprises 2,513 entries, yielding a total of 5,286 transition state configurations.

### Model training and evaluation

We implemented the QuantaMind model using the PyTorch framework.^61^ The model architecture consists of three main components: an embedding module, a stack of interaction layers, and a readout module.

The embedding module encodes system-level and atomic-level features into machine-readable vectors. These features include atomic numbers, the total system charge, spin multiplicity, and the chosen DFT functional and basis set.

The encoded information is then passed through three interaction layers, each composed of a local and a non-local interaction sublayer. The local interaction sublayer is constructed using a message-passing neural network (MPNN) framework, ^62^ inspired by recent advances in 3D molecular representation learning.^41,63^ This component captures short-range atomic interactions based on molecular geometry. In parallel, the non-local interaction sublayer, implemented using a transformer-based attention mechanism,^64^ is designed to capture long-range correlations within large biomolecular systems. Prior research has demonstrated that non-local interactions can be significant for accurate prediction in some systems.^65^

The atomic features obtained from the interaction layers are then passed to the readout module, which predicts atomic-level energy and partial charge values. The total energy of the system is computed as the sum of per-atom energies, and atomic charges are used to calculate molecular dipole moments. Atomic forces are obtained by computing the gradient of the total energy with respect to atomic coordinates.

Model training is carried out using the PyTorch Lightning framework.^66^ The learning objective is to minimize the root mean squared error (RMSE) of energy, force, and dipole moment predictions. The model is trained with the AMSGrad variant of the Adam optimizer,^67^ using a starting learning rate of 0.001. If the evaluation loss does not improve within 25 epochs, the learning rate is reduced by a factor of 10. To ensure balanced training across heterogeneous datasets, an oversampling strategy is used to equalize the chance of sampling data points from different sources during training. Additionally, Stochastic Weight Averaging (SWA)^68^ is employed to improve generalization.

In addition to the datasets developed in this work, we incorporate the general protein fragments dataset and the crambin dataset from a previous study,^16^ resulting in a total of six datasets used for model training. For dataset partitioning, an 80/10/10 split is applied to the smaller datasets, including the phosphorus dataset, the transition state dataset, and the crambin dataset. For the larger general protein fragment dataset, we use fixed splits of 5,000 data points each for evaluation and testing, with the remainder used for training. All datasets are randomly partitioned into training, validation, and test sets.

### Ab initio molecular dynamic simulation with QuantaMind

Once trained, the QuantaMind model can be used to perform molecular dynamics (MD) simulations by leveraging its predicted atomic forces. Simulations are carried out using the Atomic Simulation Environment (ASE)^69^ in conjunction with the SchNetPack framework.^70^ For simulations involving water dissociation, we use the SPC216 water box,^71^ a pre-equilibrated configuration commonly applied in GROMACS simulations. The system is evolved under periodic boundary conditions within the NVT ensemble using a 0.5 fs time step.

To model a proton transfer process, we manually modify one of the water molecules in the SPC216 box by adding an excess proton to form a hydronium ion (H3O+). For simulating acid-base neutralization, a second water molecule is selected approximately 10 Å away from the hydronium ion and modified to form a hydroxide ion (OH-). Simulations involving spontaneous water dissociation are conducted at 393 K, while simulations of proton diffusion and neutralization reactions are performed at 300 K.

Hydronium and hydroxide ions are identified in the simulation trajectory based on local hydrogen-oxygen distances. Specifically, an oxygen atom is classified as belonging to a hydronium ion if it is bonded to three hydrogen atoms within a 1.2 Å cutoff. Similarly, an oxygen atom is considered part of a hydroxide ion if it has only one hydrogen within the 1.2 Å cutoff. This threshold is chosen empirically, balancing the covalent O-H bond length in neutral water (0.958 Å) and the typical hydrogen bond length (1.97 Å).

### Structure optimization and pocket-centric simulations with QuantaMind

QuantaMind is also capable of performing energy minimization, which is implemented using the Atomic Simulation Environment (ASE).^69^ At each minimization step, QuantaMind predicts the total system energy and atomic forces based on the current geometry. The atomic coordinates are then updated using the Limited-memory Broyden-Fletcher-Goldfarb-Shanno (L-BFGS) optimizer.^72,73^ This cycle of energy and force evaluation followed by coordinate updates is repeated until the convergence criterion is met or the maximum number of optimization steps is reached.

For protein structure optimization tasks, we apply this energy minimization procedure to full protein structures, with a maximum of 100 optimization steps per structure.

For protein-ligand interaction prediction tasks, simulating the entire protein is often computationally inefficient, especially when only the ligand-bound binding pocket is of interest. To address this, we implement a pocket-centric optimization scheme based on spatial proximity to the ligand. Specifically, we define inner and outer shell cutoff distances: atoms within the inner shell radius are fully modeled and allowed to relax during optimization; atoms between the inner and outer shells are modeled but their positions remain fixed; atoms beyond the outer shell radius are excluded entirely from the simulation. This hierarchical strategy significantly reduces computational cost by restricting calculations to the relevant local environment around the ligand.

We use an inner shell cutoff of 5 Å and an outer shell cutoff of 45 Å for our pocket-centric simulations in this work.

### Protein evaluation benchmark and metrics

The evaluation benchmark set is constructed based on the OOD60 and Domain Motion datasets from the BioEmu study.^25^ The OOD60 dataset comprises 19 protein examples collected from the Protein Data Bank (PDB) after the AlphaFold2 monomer model training cut-off date (April 30, 2018). A 60% sequence identity cutoff is applied to exclude any proteins with high similarity to chains present in the PDB before the specified cutoff. The Domain Motion dataset consists of 22 protein structures that exhibit large-scale conformational changes due to hinge motions.

To reduce computational cost, we selected a subset of these datasets for evaluation. Specifically, 10 proteins were chosen from OOD60 (A0A0H3AFX3 E1C7U0 J7QA90 K0JNC6 P0A8W0 P0CL43 P21926 P50405 Q31PX7 Q9NR30), and 7 proteins from the Domain Motion dataset (B0F0C5 P03047 P06766 P0A8W0 P67701 P71447 Q18A65).

For structural accuracy assessment, we compute the fraction of native contacts, *F*_nat_, which measures the proportion of native residue-residue contacts preserved in the predicted structure. To calculate this, we identify all residue pairs within 3 Å of each other in the crystal structure, excluding sequential neighbors. These pairs are then compared against the predicted structure, and *F*_nat_ is defined as the fraction of these native contacts retained.

In addition, we evaluate side-chain hydrogen bond recovery using the side-chain hydrogen bond recovery score, *F*_h-bond_. This metric represents the proportion of native side-chain hydrogen bonds present in the crystal structure that are also observed in the predicted structure. Hydrogen bonds are identified using the MDTraj library.^74^

### Protein-ligand evaluation benchmark and metrics

We construct our protein-ligand evaluation benchmark based on the PoseBuster benchmark dataset.^35^ Most ligands in PoseBuster are chemically distinct from those present in our training data. To quantify this and ensure a fair evaluation, we compute the chemical similarity between each ligand and the 20 standard amino acids. A similarity threshold of 0.44, determined from Figure S1D, is used based on prior results indicating that ligands with similarity greater than 0.44 yield a 100% success rate. Applying this filter to the 308 protein-ligand pairs in the PoseBuster benchmark, we identify 8 examples that satisfy the similarity criterion and use these for downstream evaluation.

In addition to the 18 standard metrics included in PoseBuster, we incorporate two supplementary measures: (i) ligand RMSD, which denotes the percentage of predicted ligands with a root mean squared deviation below 2 Å relative to their respective crystal structures, and (ii) *F*_h-bond_ which quantifies the fraction of native protein-ligand hydrogen bonds recovered in the predicted complex. Hydrogen bonds and other non-covalent interactions are identified using the PLIP Python package,^75^ which is specifically designed for protein-ligand interaction profiling.

A subset of five representative PoseBuster metrics is shown in Figure 7B, while performance across all 18 metrics is provided in the supplementary Figure S2.

## Data Availability

The source code used in this study is available at https://github.com/MoleculeMindOpenSource/QuantaMind. The molecular dynamics trajectories generated in this work are accessible via Zenodo at https://zenodo.org/records/16910424.

## Author Contributions

Conceptualization: Deqiang Zhang, Jinbo Xu; Methodology: Song Xia; Software: Song Xia; MD Simulations: Xu Shang. Formal analysis: Song Xia, Deqiang Zhang; Writing - original draft: Song Xia; Writing - review & editing: Deqiang Zhang, Jinbo Xu, Song Xia; Supervision: Deqiang Zhang, Jinbo Xu; Funding acquisition: Jinbo Xu.

## Supporting Information Available

**Figure S1:**
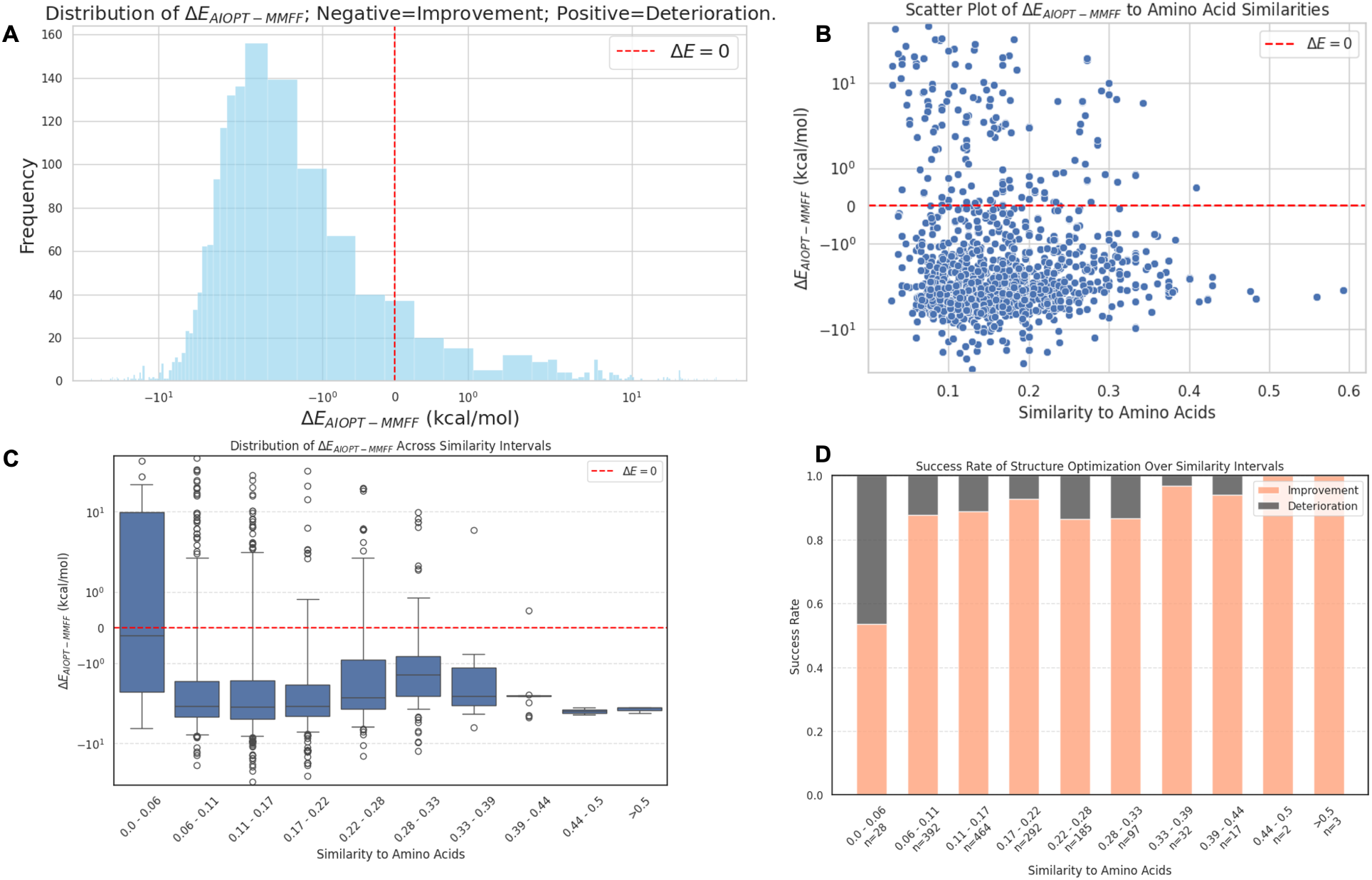
QuantaMind performance on small molecule optimization tasks.

**Figure S2:**
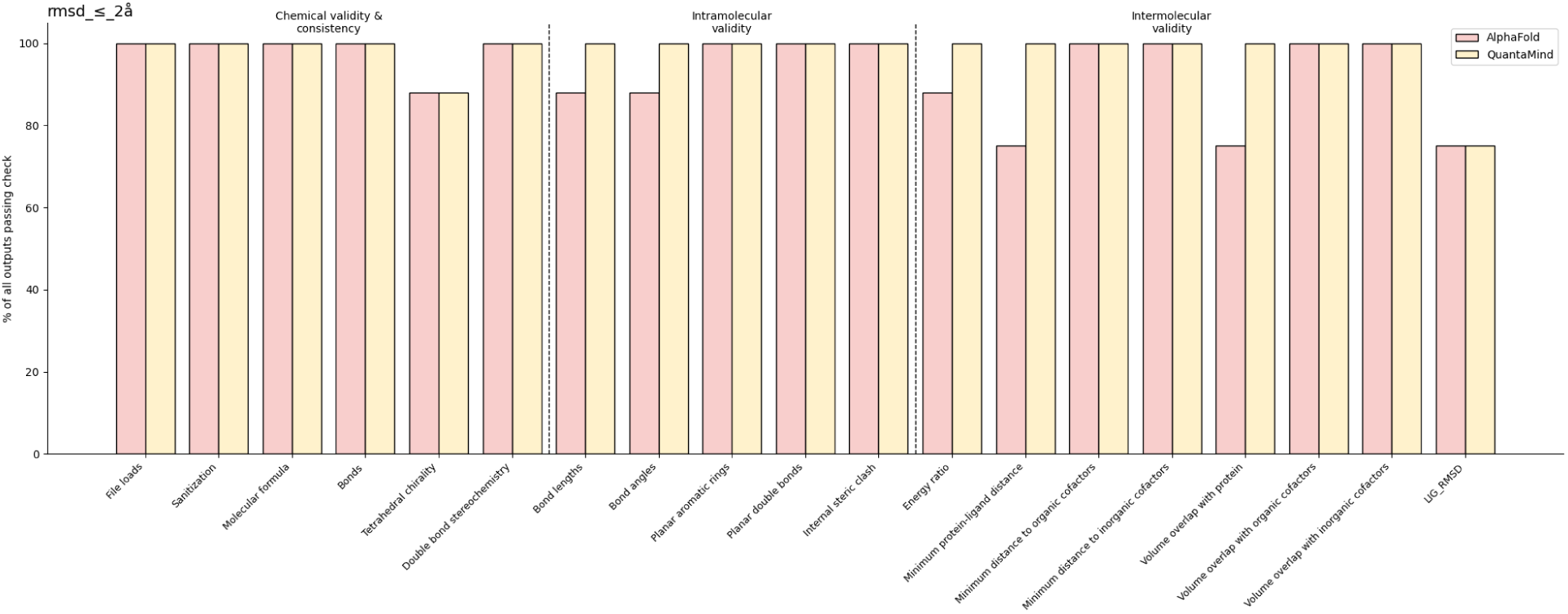
Full PoseBuster benchmark performance across all 18 evaluation metrics.

## References

(1) Hollingsworth, S. A.; Dror, R. O. Molecular dynamics simulation for all. Neuron 2018, 99, 1129–1143.

(2) Hansson, T.; Oostenbrink, C.; van Gunsteren, W. Molecular dynamics simulations. Current opinion in structural biology 2002, 12, 190–196.

(3) Karplus, M.; Petsko, G. A. Molecular dynamics simulations in biology. Nature 1990, 347, 631–639.

(4) Karplus, M.; McCammon, J. A. Molecular dynamics simulations of biomolecules. Nature structural biology 2002, 9, 646–652.

(5) Dirac, P. A. M. Quantum mechanics of many-electron systems. Proceedings of the Royal Society of London. Series A, Containing Papers of a Mathematical and Physical Character 1929, 123, 714–733.

(6) Gordon, M. S.; Schmidt, M. W. Theory and applications of computational chemistry; Elsevier, 2005; pp 1167–1189.

(7) Iftimie, R.; Minary, P.; Tuckerman, M. E. Ab initio molecular dynamics: Concepts, recent developments, and future trends. Proceedings of the National Academy of Sciences 2005, 102, 6654–6659.

(8) Halgren, T. A. Merck molecular force field. II. MMFF94 van der Waals and electrostatic parameters for intermolecular interactions. Journal of Computational Chemistry 1996, 17, 520–552.

(9) González, M. A. Force fields and molecular dynamics simulations. École thématique de la Société Française de la Neutronique 2011, 12, 169–200.

(10) Unke, O. T.; Chmiela, S.; Sauceda, H. E.; Gastegger, M.; Poltavsky, I.; Schutt, K. T.; Tkatchenko, A.; Muller, K.-R. Machine learning force fields. Chemical Reviews 2021, 121, 10142–10186.

(11) Langer, M. F.; Goeßmann, A.; Rupp, M. Representations of molecules and materials for interpolation of quantum-mechanical simulations via machine learning. npj Computational Materials 2022, 8, 41.

(12) Unke, O. T.; Koner, D.; Patra, S.; Käser, S.; Meuwly, M. High-dimensional potential energy surfaces for molecular simulations: from empiricism to machine learning. Machine Learning: Science and Technology 2020, 1, 013001.

(13) Unke, O. T.; Meuwly, M. PhysNet: A neural network for predicting energies, forces, dipole moments, and partial charges. Journal of chemical theory and computation 2019, 15, 3678–3693.

(14) Klicpera, J.; Groß, J.; Günnemann, S.; others Directional message passing for molecular graphs. ICLR. 2020; pp 1–13.

(15) Wang, Y.; Wang, T.; Li, S.; He, X.; Li, M.; Wang, Z.; Zheng, N.; Shao, B.; Liu, T.-Y. Enhancing geometric representations for molecules with equivariant vector-scalar interactive message passing. Nature Communications 2024, 15, 313.

(16) Unke, O. T.; Stöhr, M.; Ganscha, S.; Unterthiner, T.; Maennel, H.; Kashubin, S.; Ahlin, D.; Gastegger, M.; Sandonas, L. M.; Berryman, J. T.; Tkatchenko, A.; Müller, K.-R. Biomolecular dynamics with machine-learned quantum-mechanical force fields trained on diverse chemical fragments. Science Advances 2024, 10, eadn4397.

(17) Wang, T.; He, X.; Li, M.; Li, Y.; Bi, R.; Wang, Y.; Cheng, C.; Shen, X.; Meng, J.; Zhang, H.; others Ab initio characterization of protein molecular dynamics with AI2BMD. Nature 2024, 1–9.

(18) Schlick, T.; Portillo-Ledesma, S. Biomolecular modeling thrives in the age of technology. Nature computational science 2021, 1, 321–331.

(19) Teale, A. M.; Helgaker, T.; Savin, A.; Adamo, C.; Aradi, B.; Arbuznikov, A. V.; Ayers, P. W.; Baerends, E. J.; Barone, V.; Calaminici, P.; others DFT exchange: sharing perspectives on the workhorse of quantum chemistry and materials science. Physical chemistry chemical physics 2022, 24, 28700–28781.

(20) Bursch, M.; Mewes, J.-M.; Hansen, A.; Grimme, S. Best-practice DFT protocols for basic molecular computational chemistry. Angewandte Chemie 2022, 134, e202205735.

(21) Mohammed, R.; Rawashdeh, J.; Abdullah, M. Machine learning with oversampling and undersampling techniques: overview study and experimental results. 2020 11th international conference on information and communication systems (ICICS). 2020; pp 243–248.

(22) Einstein, A. UN THE MOVEMENT OF SMALL PARTICLES SUSPENDED IN STATIUNARY LIQUIDS REQUIRED BY THE MOLECULAR-KINETIC THEORY 0F HEAT. 1905,

(23) Tuckerman, M. E. Statistical mechanics: theory and molecular simulation; Oxford university press, 2023.

(24) Atkins, P. W.; De Paula, J.; Keeler, J. Atkins’ physical chemistry; Oxford university press, 2023.

(25) Lewis, S.; Hempel, T.; Jiménez-Luna, J.; Gastegger, M.; Xie, Y.; Foong, A. Y.; Satorras, V. G.; Abdin, O.; Veeling, B. S.; Zaporozhets, I.; others Scalable emulation of protein equilibrium ensembles with generative deep learning. Science 2025, eadv9817.

(26) Lu, J.; Zhong, B.; Zhang, Z.; Tang, J. Str2str: A score-based framework for zero-shot protein conformation sampling. arXiv preprint arXiv:2306.03117 2023,

(27) Lu, J.; Chen, X.; Lu, S. Z.; Shi, C.; Guo, H.; Bengio, Y.; Tang, J. Structure Language Models for Protein Conformation Generation. arXiv preprint arXiv:2410.18403 2024,

(28) Jing, B.; Berger, B.; Jaakkola, T. AlphaFold Meets Flow Matching for Generating Protein Ensembles. Forty-first International Conference on Machine Learning. 2024.

(29) Visani, G. M.; Galvin, W.; Pun, M.; Nourmohammad, A. H-Packer: Holographic Rotationally Equivariant Convolutional Neural Network for Protein Side-Chain Packing. Machine Learning in Computational Biology. 2024; pp 230–249.

(30) McPartlon, M.; Xu, J. An end-to-end deep learning method for protein side-chain packing and inverse folding. Proceedings of the National Academy of Sciences 2023, 120, e2216438120.

(31) Das, R.; Baker, D. Macromolecular modeling with rosetta. Annu. Rev. Biochem. 2008, 77, 363–382.

(32) Mirdita, M.; Schütze, K.; Moriwaki, Y.; Heo, L.; Ovchinnikov, S.; Steinegger, M. ColabFold: making protein folding accessible to all. Nature methods 2022, 19, 679–682.

(33) Jumper, J.; Evans, R.; Pritzel, A.; Green, T.; Figurnov, M.; Ronneberger, O.; Tunyasuvunakool, K.; Bates, R.; Žídek, A.; Potapenko, A.; others Highly accurate protein structure prediction with AlphaFold. nature 2021, 596, 583–589.

(34) Abramson, J.; Adler, J.; Dunger, J.; Evans, R.; Green, T.; Pritzel, A.; Ronneberger, O.; Willmore, L.; Ballard, A. J.; Bambrick, J.; others Accurate structure prediction of biomolecular interactions with AlphaFold 3. Nature 2024, 630, 493–500.

(35) Buttenschoen, M.; Morris, G. M.; Deane, C. M. PoseBusters: AI-based docking methods fail to generate physically valid poses or generalise to novel sequences. Chemical Science 2024, 15, 3130–3139.

(36) Abramson, J. et al. Accurate structure prediction of biomolecular interactions with AlphaFold 3. Nature 2024, 630, 493–500.

(37) Reymond, J.-L. The chemical space project. Accounts of chemical research 2015, 48, 722–730.

(38) Xia, S.; Chen, E.; Zhang, Y. Integrated molecular modeling and machine learning for drug design. Journal of chemical theory and computation 2023, 19, 7478–7495.

(39) von Lilienfeld, O. A.; Müller, K.-R.; Tkatchenko, A. Exploring chemical compound space with quantum-based machine learning. Nature Reviews Chemistry 2020, 4, 347– 358.

(40) Smith, J. S.; Nebgen, B.; Lubbers, N.; Isayev, O.; Roitberg, A. E. Less is more: Sampling chemical space with active learning. The Journal of chemical physics 2018, 148.

(41) Lu, J.; Xia, S.; Lu, J.; Zhang, Y. Dataset Construction to Explore Chemical Space with 3D Geometry and Deep Learning. Journal of Chemical Information and Modeling 2021, 61, 1095–1104.

(42) Smith, J. S.; Isayev, O.; Roitberg, A. E. ANI-1, A data set of 20 million calculated off-equilibrium conformations for organic molecules. Scientific data 2017, 4, 1–8.

(43) Chanussot, L.; Das, A.; Goyal, S.; Lavril, T.; Shuaibi, M.; Riviere, M.; Tran, K.; Heras-Domingo, J.; Ho, C.; Hu, W.; others Open catalyst 2020 (OC20) dataset and community challenges. Acs Catalysis 2021, 11, 6059–6072.

(44) Ramakrishnan, R.; Dral, P. O.; Rupp, M.; Von Lilienfeld, O. A. Quantum chemistry structures and properties of 134 kilo molecules. Scientific data 2014, 1, 1–7.

(45) Schreiner, M.; Bhowmik, A.; Vegge, T.; Busk, J.; Winther, O. Transition1x-a dataset for building generalizable reactive machine learning potentials. Scientific Data 2022, 9, 779.

(46) Anstine, D. M.; Zhao, Q.; Zubatiuk, R.; Zhang, S.; Singla, V.; Nikitin, F.; Savoie, B. M.; Isayev, O. AIMNet2-rxn: A Machine Learned Potential for Generalized Reaction Modeling on a Millions-of-Pathways Scale. 2025,

(47) Yuan, E. C.-Y.; Kumar, A.; Guan, X.; Hermes, E. D.; Rosen, A. S.; Zádor, J.; Head-Gordon, T.; Blau, S. M. Analytical ab initio hessian from a deep learning potential for transition state optimization. Nature Communications 2024, 15, 8865.

(48) Hermes, E. D.; Sargsyan, K.; Najm, H. N.; Zádor, J. Sella, an open-source automation-friendly molecular saddle point optimizer. Journal of Chemical Theory and Computation 2022, 18, 6974–6988.

(49) Hermes, E. D.; Sargsyan, K.; Najm, H. N.; Zádor, J. Accelerated saddle point refinement through full exploitation of partial hessian diagonalization. Journal of chemical theory and computation 2019, 15, 6536–6549.

(50) Hermes, E. D.; Sargsyan, K.; Najm, H. N.; Zádor, J. Geometry optimization speedup through a geodesic approach to internal coordinates. The Journal of Chemical Physics 2021, 155.

(51) Isiorho, E. A.; Jeon, B.-S.; Kim, N. H.; Liu, H.-w.; Keatinge-Clay, A. T. Structural studies of the spinosyn forosaminyltransferase, SpnP. Biochemistry 2014, 53, 4292– 4301.

(52) Trott, O.; Olson, A. J. AutoDock Vina: improving the speed and accuracy of docking with a new scoring function, efficient optimization, and multithreading. Journal of computational chemistry 2010, 31, 455–461.

(53) Parrish, R. M.; Burns, L. A.; Smith, D. G.; Simmonett, A. C.; DePrince III, A. E.; Hohenstein, E. G.; Bozkaya, U.; Sokolov, A. Y.; Di Remigio, R.; Richard, R. M.; others Psi4 1.1: An open-source electronic structure program emphasizing automation, advanced libraries, and interoperability. Journal of chemical theory and computation 2017, 13, 3185–3197.

(54) Adamo, C.; Barone, V. Toward reliable density functional methods without adjustable parameters: The PBE0 model. The Journal of chemical physics 1999, 110, 6158–6170.

(55) Tkatchenko, A.; DiStasio Jr, R. A.; Car, R.; Scheffler, M. Accurate and efficient method for many-body van der Waals interactions. Physical review letters 2012, 108, 236402.

(56) Ahmadi, S.; Barrios Herrera, L.; Chehelamirani, M.; Hostaš, J.; Jalife, S.; Salahub, D. R. Multiscale modeling of enzymes: QM-cluster, QM/MM, and QM/MM/MD: a tutorial review. International Journal of Quantum Chemistry 2018, 118, e25558.

(57) Camm, E. L.; Towers, G. N. Phenylalanine ammonia lyase. Phytochemistry 1973, 12, 961–973.

(58) Chan, H.-C.; Zhu, Y.; Hu, Y.; Ko, T.-P.; Huang, C.-H.; Ren, F.; Chen, C.-C.; Ma, Y.; Guo, R.-T.; Sun, Y. Crystal structures of D-psicose 3-epimerase from Clostridium cellulolyticum H10 and its complex with ketohexose sugars. Protein & cell 2012, 3, 123–131.

(59) Ferrandi, E. E.; Sayer, C.; Isupov, M. N.; Annovazzi, C.; Marchesi, C.; Iacobone, G.; Peng, X.; Bonch-Osmolovskaya, E.; Wohlgemuth, R.; Littlechild, J. A.; others Discovery and characterization of thermophilic limonene-1, 2-epoxide hydrolases from hot spring metagenomic libraries. The FEBS Journal 2015, 282, 2879–2894.

(60) Frisch, M. J. et al. Gaussian16 Revision C.01. 2016; Gaussian Inc. Wallingford CT.

(61) Paszke, A. Pytorch: An imperative style, high-performance deep learning library. arXiv preprint arXiv:1912.01703 2019,

(62) Gilmer, J.; Schoenholz, S. S.; Riley, P. F.; Vinyals, O.; Dahl, G. E. Neural message passing for quantum chemistry. International conference on machine learning. 2017; pp 1263–1272.

(63) Xia, S.; Zhang, D.; Zhang, Y. Multitask Deep Ensemble Prediction of Molecular Energetics in Solution: From Quantum Mechanics to Experimental Properties. Journal of Chemical Theory and Computation 2023, 19, 659–668.

(64) Vaswani, A.; Shazeer, N.; Parmar, N.; Uszkoreit, J.; Jones, L.; Gomez, A. N.; Kaiser, Ł.; Polosukhin, I. Attention is all you need. Advances in neural information processing systems 2017, 30.

(65) Unke, O. T.; Chmiela, S.; Gastegger, M.; Schütt, K. T.; Sauceda, H. E.; Müller, K.-R. SpookyNet: Learning force fields with electronic degrees of freedom and nonlocal effects. Nature communications 2021, 12, 7273.

(66) Falcon, W.; The PyTorch Lightning team PyTorch Lightning. 2019; https://github.com/Lightning-AI/lightning.

(67) Reddi, S. J.; Kale, S.; Kumar, S. On the convergence of adam and beyond. arXiv preprint arXiv:1904.09237 2019,

(68) Izmailov, P.; Podoprikhin, D.; Garipov, T.; Vetrov, D.; Wilson, A. G. Averaging weights leads to wider optima and better generalization. arXiv preprint arXiv:1803.05407 2018,

(69) Larsen, A. H.; Mortensen, J. J.; Blomqvist, J.; Castelli, I. E.; Christensen, R.; Dułak, M.; Friis, J.; Groves, M. N.; Hammer, B.; Hargus, C.; others The atomic simulation environmenta Python library for working with atoms. Journal of Physics: Condensed Matter 2017, 29, 273002.

(70) Schutt, K.; Kessel, P.; Gastegger, M.; Nicoli, K. A.; Tkatchenko, A.; Muller, K.-R. SchNetPack: A deep learning toolbox for atomistic systems. Journal of chemical theory and computation 2018, 15, 448–455.

(71) Berendsen, H. J.; Postma, J. P.; van Gunsteren, W. F.; Hermans, J. Interaction models for water in relation to protein hydration. Intermolecular forces: proceedings of the fourteenth Jerusalem symposium on quantum chemistry and biochemistry held in jerusalem, israel, april 13–16, 1981. 1981; pp 331–342.

(72) Liu, D. C.; Nocedal, J. On the limited memory BFGS method for large scale optimization. Mathematical programming 1989, 45, 503–528.

(73) Fletcher, R. Practical methods of optimization; John Wiley & Sons, 2000.

(74) McGibbon, R. T.; Beauchamp, K. A.; Harrigan, M. P.; Klein, C.; Swails, J. M.; Hernández, C. X.; Schwantes, C. R.; Wang, L.-P.; Lane, T. J.; Pande, V. S. MDTraj: A Modern Open Library for the Analysis of Molecular Dynamics Trajectories. Biophysical Journal 2015, 109, 1528 – 532.

(75) Adasme, M. F.; Linnemann, K. L.; Bolz, S. N.; Kaiser, F.; Salentin, S.; Haupt, V. J.; Schroeder, M. PLIP 2021: expanding the scope of the protein–ligand interaction profiler to DNA and RNA. Nucleic acids research 2021, *49*, W530–W534.

